# Revised International Staging System (R-ISS) stage-dependent analysis uncovers oncogenes and potential immunotherapeutic targets in multiple myeloma

**DOI:** 10.1101/2021.12.06.471423

**Authors:** Ling Zhong, Xinwei Yuan, Qian Zhang, Tao Jiang, Huan Li, Jialing Xiao, Chenglong Li, Lan Luo, Ping Shuai, Liang Wang, Yuping Liu, Man Yu, Yi Shi, Wei Zhang, Yunbin Zhang, Bo Gong

## Abstract

Multiple myeloma (MM), characterized by high intratumour heterogeneity, accounts for ∼10% of all haematologic malignancies. Stratified by the Revised International Staging System (R-ISS), little is known about R-ISS-related plasma cell (PC) heterogeneity, gene expression modules in cytotoxic T/NK cells and immunoregulatory ligands and receptors. Herein, we constructed a single-cell transcriptome atlas of bone marrow in normal and R-ISS-staged MM patients. Focusing on PCs, we identified and validated a subset of GZMA+ cytotoxic PCs. In addition, a malignant PC population with high proliferation capability (proliferating PCs) was associated with unfavourable prognosis and EBV infection in our collected samples. Ribonucleotide Reductase Regulatory Subunit M2 (RRM2), a specific marker of proliferating PCs, was shown to induce MM cell line proliferation and serve as a detrimental marker in MM. Subsequently, three R-ISS-dependent gene modules in cytotoxic CD8+ T and NKT cells were identified and functionally analysed. Finally, cell-cell communication between neutrophils and proliferating PCs with cytotoxic CD8+ T and NKT cells was investigated, which identified intercellular ligand receptors and potential immunotargets such as SIRPA-CD47 and TIGIT-NECTIN3. Collectively, this study provides an R-ISS-related single-cell MM atlas and reveals the clinical significance of two PC clusters, as well as potential immunotargets in MM progression.

## Introduction

Multiple myeloma (MM) is characterized by uncontrolled proliferation of monoclonal plasma cells and accounts for ∼10% of all haematologic malignancies [1]. The revised International Staging System (R-ISS) was developed to stratify MM patients into groups I, II and III [2] with distinct outcomes [3] and treatment response [4], with or without the assistance of other clinical parameters [5, 6]. However, the single-cell gene expression signatures in MM R-ISS stages remain to be explored. Epstein-Barr virus (EBV), the first human tumour virus [7], causes Burkitt, Hodgkin, and post-transplant B cell lymphomas [8], and is associated with poor prognosis and clinical characteristics of RISS III in MM patients [9], although further studies are needed to uncover the mechanism. In the bone marrow (BM) microenvironment, the interplay between neoplastic cells and immune microenvironment cells is involved in MM progression and drug response [10], and single-cell level ligand-receptor interactions remain unclear. Myeloid cells (such as neutrophils) foster cancer-promoting inflammation, and natural killer (NK) cells and T lymphocytes mediate protective antitumour responses [11]. With dramatic advances in immunomodulatory drugs [12-14], monoclonal antibodies [15], proteasome inhibitors [16], and histone deacetylase inhibitors (HDACis) [17], MM patients still remain largely incurable [18].

Single-cell sequencing (ScRNA-Seq) offers an unprecedented opportunity to study the heterogeneity of plasma cells and immune microenvironments in cancer. Focusing on plasma cells, intratumour heterogeneity (ITH) [19], genome evolution [20] and transcriptome expression signatures [21], resistance pathways and therapeutic targets in relapsed MM [22] were revealed. Meanwhile, compromised microenvironment immune cells [23] and transcriptional alterations [24] in MM precursor stages and extramedullary progression were recently uncovered. Nevertheless, little is known about malignant plasma cell and immune cell gene expression signatures with respect to the R-ISS stage and their role in EBV-infected MM at the single-cell level.

Herein, we adopted single-cell transcriptome sequencing to investigate the gene expression profiles in the normal and R-ISS stage I, II and III groups. First, we examined the heterogeneity of plasma cells and validated the function and clinical significance of two rare plasma cell populations. The function and clinical significance of proliferating PCs and the hub gene RRM2 were validated in other cohorts, cell lines and collected samples. Subsequently, gene expression modules underlying two T cell clusters with decreased proportions along with MM R-ISS stage were investigated. Finally, cell-cell communication was analysed to interpret the tumour cell-cytotoxic T cell and cytotoxic T cell-neutrophil interactions in MM. Collectively, the results of this study provide an R-ISS-related single-cell MM atlas and reveal the clinical significance of two PC clusters, as well as potential immunotargets in MM progression.

## Results

### Single-cell transcriptome atlas of R-ISS-staged multiple myeloma

To explore the intratumoural heterogeneity of R-ISS stage-classified multiple myeloma, 11 bone marrow samples from 2 healthy donors (healthy control, HC), 2 R-ISS I, 2 R-ISS II and 5 R-ISS III stage MM patients were subjected to single-cell suspension parathion and transcriptome sequencing (**Fig 1A**). After low-quality cell filtering and quality control, 103,043 single-cell expression matrixes were acquired. Subsequent dimensional reduction generated 21 clusters (**Fig 1B**). Most clusters existed in all four groups (**Fig 1C**), showing a batch-effect removal basis for the following analysis. Then, based on the cell type markers, six general cell types were identified, including plasma cells, B cells, myeloid cells, CD4+ T cells, CD8+ T cells and immature red cells (**Fig 1D-1E)**. The proportions of 21 cell clusters in all groups are shown in **Fig 1F**.

**Figure 1.**
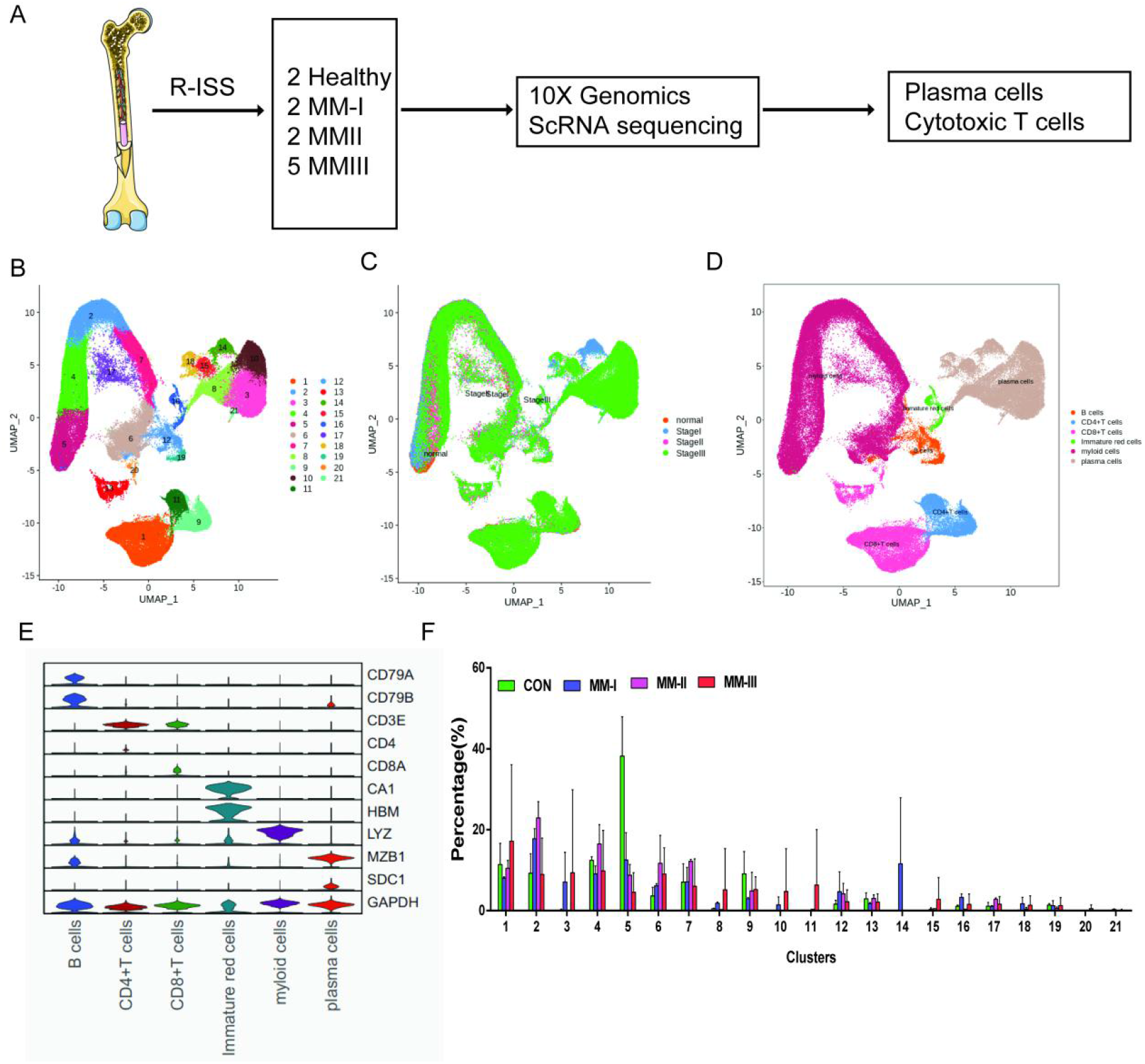
Single cell transcriptome atlas of Multiple myeloma (MM) with R-ISS staging. A) Schematic illustration of workflow in this study; B) Dimension reduction of cells, and 21 clusters were acquired, shown with UMAP; C) UMAP showing the distribution of sample groups of Normal, R-ISS I to III; Based on expression signature of canonical markers, six general cell types were identified. UMAP of cell type in were shown D), and violin plot of cell type markers in E); F) Proportion of cell clusters in normal and MM RISS groups;

### Functional identification of a rare cytotoxic NKG7+GZMA+plasma cell population in MM

Intraclonal heterogeneity in plasma cells is emerging as a vital modulator in MM progression [25], drug sensitivity and therapeutic response [26]. In **Fig 1D**, 7 clusters (3, 8, 10, 14, 15, 18 and 21) specifically expressing high levels of CD138 were classified into plasma cells. Then, specific markers in each plasma cell cluster were calculated. All 7 clusters showed abundant expression of ribosomal proteins such as RPS2 and RPL26, which corresponds to the antibody-producing function of plasma cells. Based on the expression of specific markers, all 7 clusters were generally classified into two groups, group 1: C3, C8, C10, C21 and group 2: C14, C15, and C18, consistent with the layout of these clusters in Fig 1B. Comparatively, increased expression of CCL3, NES and S100A8, DEFA3, S100A9, S100A12 and CAMP was observed in group 1 and group 2, respectively (**Fig 2A**). Notably, C21 exhibits a marker expression signature distinct from other clusters in the two groups. The cytotoxic genes NKG7, GZMA, and GNLY and the chemokines CCL5 and CCL4 were exclusively expressed in C21 cells and were defined as “cytotoxic plasma cells”. Maria et al. reported that plasma cells producing granzyme B (GZMB) showed cancer cell cytotoxic activities [27], but studies of cytotoxic plasma cells in MM remain limited. The existence of cytotoxic plasma cells prompted us to investigate their existence and clinical relevance in MM. Considering that C21 accounts for a rare population (average 2.04%, ranging from 0%-10.00%) in all plasma cells (**Fig 2B**), we first validated its existence with another MM single-cell dataset. In studies focusing on plasma cell heterogeneity of symptomatic and asymptomatic myeloma (dataset GSE117156) [19], 4 markers of C21 were indeed exclusively expressed in one plasma cell cluster (cluster 9 in GSE117156, c9) (**Fig 2C-2D**). Of all 42 samples, the cell proportion varied from 0% to 30.95% of all plasma cells, with an average percentage of 4.28%, and 32 samples (76.19%) showed a percentage <5% (**Fig 2E**). In summary, we characterized a rare NKG7+GZMA+ plasma cell population with cytotoxic activity, which may provide a novel perspective for the cytotherapy of MM.

**Figure 2.**
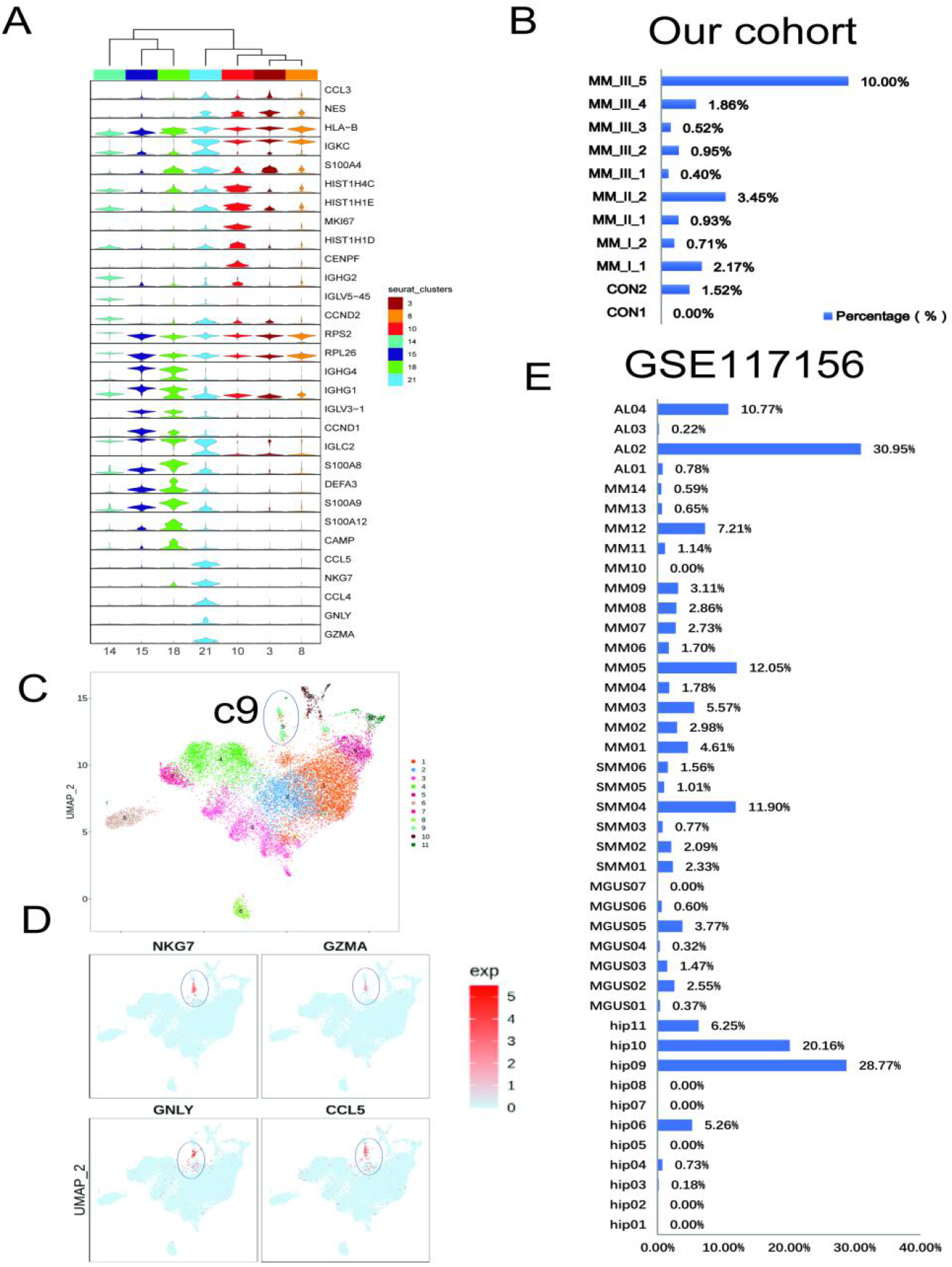
Identification of a rare cytotoxic NKG7+GZMA+plasma cell population in MM. A) The heterogeneity of plasma cells (CD138+) was transcriptionally analyzed, and genes specifically expressed in 7 plasma cell clusters were calculated. B) In our cohort, C21 accounts for a rare population (average 2.04%, ranges from 0%-10.00%) in all plasma cells; C) C21 corresponds to c9 in another MM plasma dataset of GSE117156; D) C21 specifically highly expresses cytotoxic markers like NKG7, GZMA, GNLY and CCL5; E) The cell fraction of c9 in another MM plasma dataset of GSE117156.

### Clinical significance of a malignant plasma cell population with high proliferation potential

Copy number is a common feature of MM, which usually interferes with cell cycle checkpoints to prompt accelerated proliferation [28]. To explore the oncogenic gene expression and signal transduction of plasma clusters in MM, we first conducted InferCNV to delineate the copy number variation (CNV) signals of all clusters. As shown in **Fig 3A**, strong copy number signal alterations were observed in clusters 3, 8 and 10, indicating their malignant features as tumour cells. Consistent with bulk genomic sequencing results, gain/amplification and deletion signals were mainly located on chromosomes 1, 8 and 2, 3 and 21 [29, 30]. Then we calculated the cell cycle of plasma cell clusters, and distinguished from other clusters, plasma cells in clusters 8 (C8) and 10 (C10) were presumably enriched in G2/M stage (**Fig 3B**). Expression of cell proliferation, cell cycle related oncogenic markers MKI67, TOP2A and CDK1 also showed similar pattern with overall cell cycle, suggesting a high proliferation potential of C8 and C10 in MM (**Fig 3B**). In R-ISS staged samples, C8 showed remarkably higher proportion in III stage group (**Fig1F**), and C10 were only detected in R-ISS I and III groups. Similar to C21, the existence of C10 was also validated in dataset GSE117156 (**Fig 3C**). Then GSEA analysis was conducted to dig into the signaling transduction in all 3 malignant clusters C3, C8 and C10.

**Figure 3.**
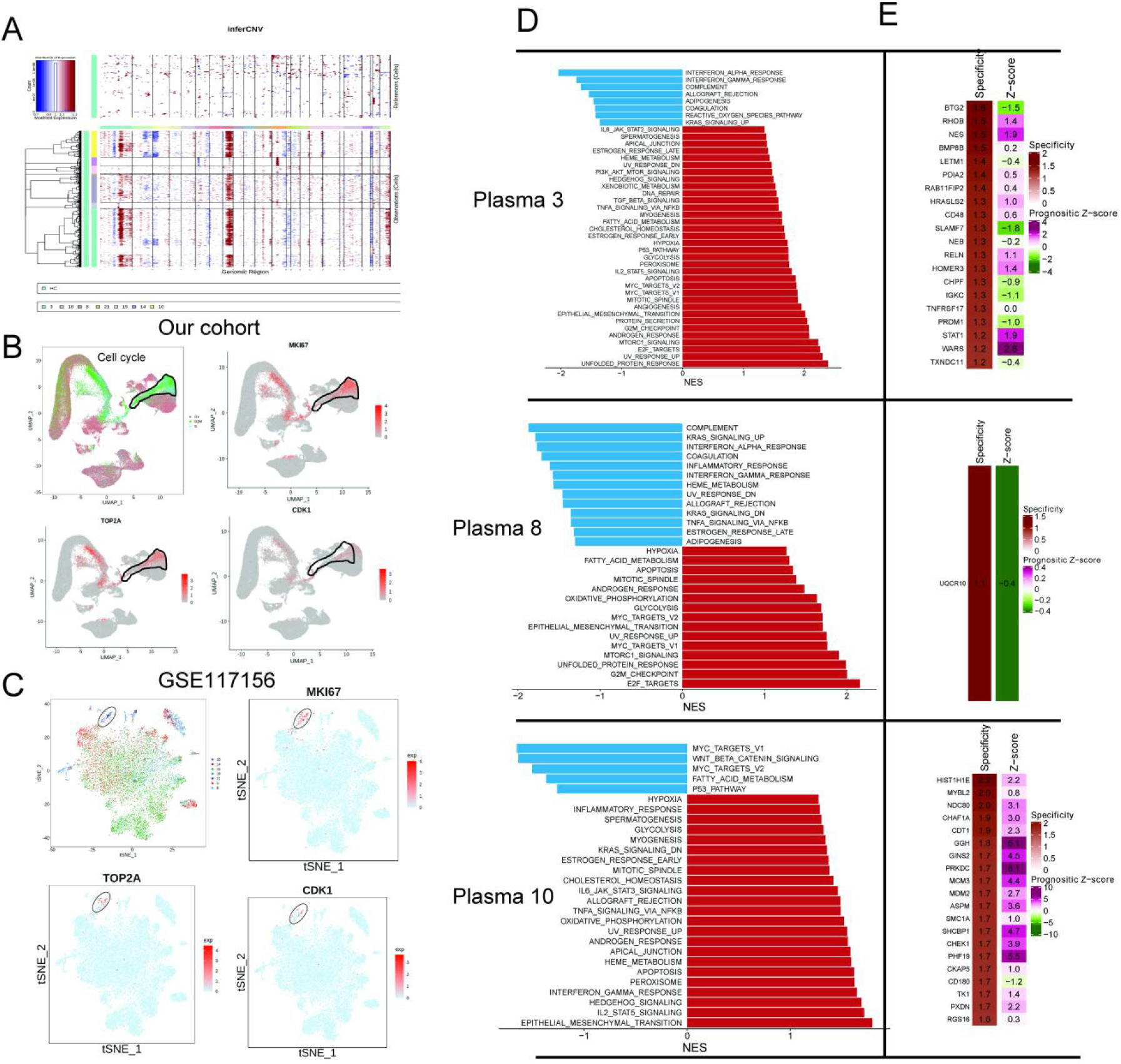
Identification of a malignant and risky plasma cell cluster with proliferation activity. A) To identify malignant plasma cells, inferCNV was applied to obtain the CNV signals in all 7 plasma cell clusters. B) Cell cycle distribution was analyzed, and shown with UMAP. UMAP of 3 proliferation markers of MKI67, TOP2A and CDK1 were also shown. C) UMAP of 3 proliferation markers of MKI67, TOP2A and CDK1 in GSE117156. D) GSEA hallmarks analysis were conducted, and significant enriched pathways of clusters 3,8 and 10 were shown. E) Precog was applied to infer the genes with specificity and prognostic Z-score in external MM gene expression data GSE6647;

As shown in **Fig 3D**, unfolded protein response, E2F targets and epithelial mesenchymal transition were the most enriched hallmark pathways in C3, C8 and C10, indicating their functional distinct roles in MM [31-34]. Finally, we applied PRECOG [35] to delineate the gene expression specificity in clusters and the prognostic score in MM with bulk sequencing GSE6477 metadata. As a result, most genes with high specificity in C10 also showed elevated prognostic Z scores, such as GGH (z-score=6.1), GINS2 (z-score=4.5), PRKDC (z-score=6.1), MCM3 (z-score=4.4), SHCBP1 (z-score=4.7), CHEK1 (z-score=3.9) and PHF19 (z-score=5.5) (**Fig 3E**). Therefore, we focused on C10, a malignant plasma cell population with high proliferation potential and unfavourable prognostic significance, and investigated its clinical relevance and potential therapeutic targets in MM.

Next, we discovered 75 significantly up-regulated genes in the MM (UGM) dataset of GSE6477 as compared with normal samples (**Fig 4A-4B**). The top 150 specific genes in C10 were compared with 75 UGMs, and 7 genes (CADM1, HIST1H1C, CD48, RRM2, PPIB, LDHB and HINT1) were acquired (**Fig 4C**). The expression of 7 UGMs is shown with the R-ISS stage in **Fig 4D**. We calculated a 7-gene signature score and analyzed the relevance of the score with respect to clinical parameters (**Fig 4E**). Next, the prognostic significance of these 7 genes was analyzed. RRM2 and HINT1 showed good performance as unfavorable markers, with HR=2.3 (95% CI=1.4-3.6, p-value<0.000402) and HR=1.9 (95% CI=1.2-2.9, p-value= 0.005496), respectively (**Fig 4F**). Then, the expression of RRM2 and HINT1 was examined in MM patients and MM.1S and U266 MM cell lines. Consistently, significantly increased expression of RRM2 and HINT1 was observed in both clinical MM R-ISS III samples (**Fig 4G**) and MM cell lines (MM.1S and U266) (**Fig 4H**). Next, the functions of RRM2 and HINT1 in the MM cell line U266 were studied. RRM2 and HINT1 silencing reduced the proliferation of U266 cells, respectively (**Fig 4I**).

**Figure 4.**
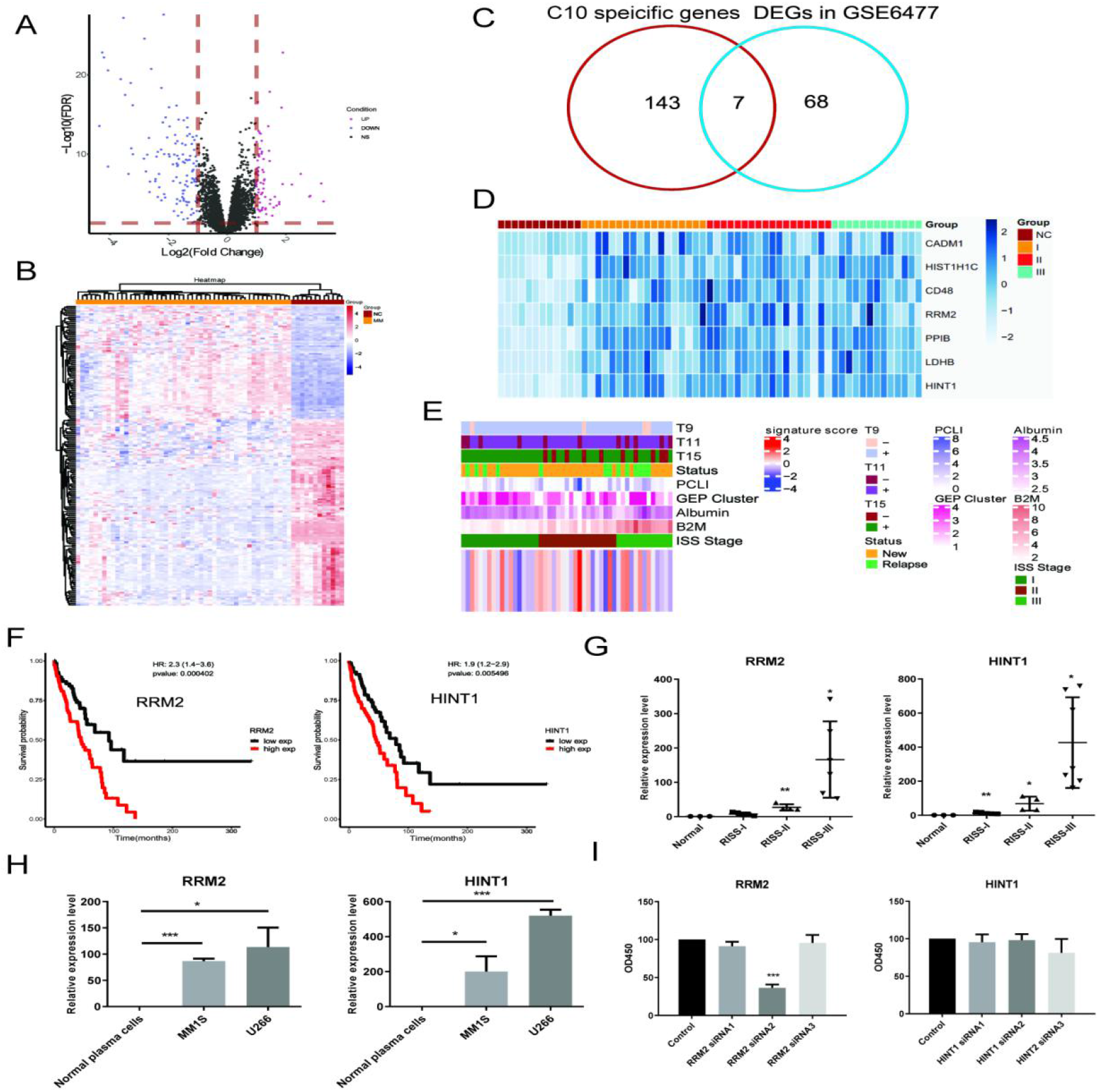
Bulk sequencing validation highlights C10 specific genes RRM2 and HINT1 as novel prognostic markers in MM. All 75 significant deregulated genes in GSE6647 dataset MM group was acquired, and shown with A) volcano plot and B) heatmap; C) Top 150 specific genes with markers potential was compared with 75 DEGs in MM dataset of GSE6647, and 7 genes were acquired and named as deregulated proliferating marker genes in MM (DPMGs); D) The expression of 7 DPMGs in normal and RISS-I to III stages were shown, and all 7 DPMGs were up-regulated in MM, especially in RISS-III stage; E) Clinical parameters of MM samples with 7-gene signature score. F) Two DPMGs, RRM2 and HINT1, exhibited good performance as unfavorable prognostic markers in MM patients. G) Relative quantification of RRM2 and HINT1 by qRT-PCR in healthy and RISS stratified MM patient BM samples; H) Relative quantification of RRM2 and HINT1 by qRT-PCR in normal plasma cells and MM1S and U266 cell lines; I) Proliferation phenotype of RRM2 and HINT1 silencing in MM cell line U266. Values represent the means of three experiments ± SD; * p < 0.05, ** p < 0.01, versus untreated control.

Finally, we calculated the differentially expressed genes (DEGs) with fold change >=2 or < =0.5 and adjusted p value <0.05 in C10 by comparing R-ISS stages I and III. All 150 DEGs were acquired. Then, we constructed a functional analysis of these 150 DEGs. As shown in **Fig 5A**, 5 functional gene modules were identified: a) ribosome, b) protein processing in endoplasmic reticulum, c) oxidative phosphorylation, d) proteasome, and e) Epstein-Barr virus infection. It is not surprising that protein and energy metabolism-related modules, such as ribosomes in protein translation, protein processing in the endoplasmic reticulum and proteasomes in protein degradation and oxidative phosphorylation, are enriched, which provides a synthetic basis for MM progression. Intriguingly, 10 genes (MDM2, CCND2, CDK6, STAT3, HLA-F, HLA-B, HLA-C, HLA-E, JUN, PSMC1) involved in Epstein-Barr virus infection and sub-modules of viral carcinogenesis attracted our attention. We then validated the expression of MKI67 and PCNA, two proliferating markers in MM patients. Indeed, significantly elevated expression of MKI67 and PCNA was observed in EBV-positive (EBV+) MM patients compared with EBV-negative (EBV-) MM patients (**Fig 5B-5C**). Finally, potential therapeutic drugs were analysed based on pathways in C10, and alsterpaullone (cyclin-dependent kinase inhibitor) [36], orlistat (anti-obesity drug) [37], moxonidine (a selective imidazoline/alpha2 adrenergic receptor agonist)[38], nalidixic acid (topoisomerase II inhibitors) [39] and LY-294002 (PI3K/AKT inhibitor) [40] were proposed as C10-targeting pharmaceuticals in MM (**Fig 5D**).

**Figure 5.**
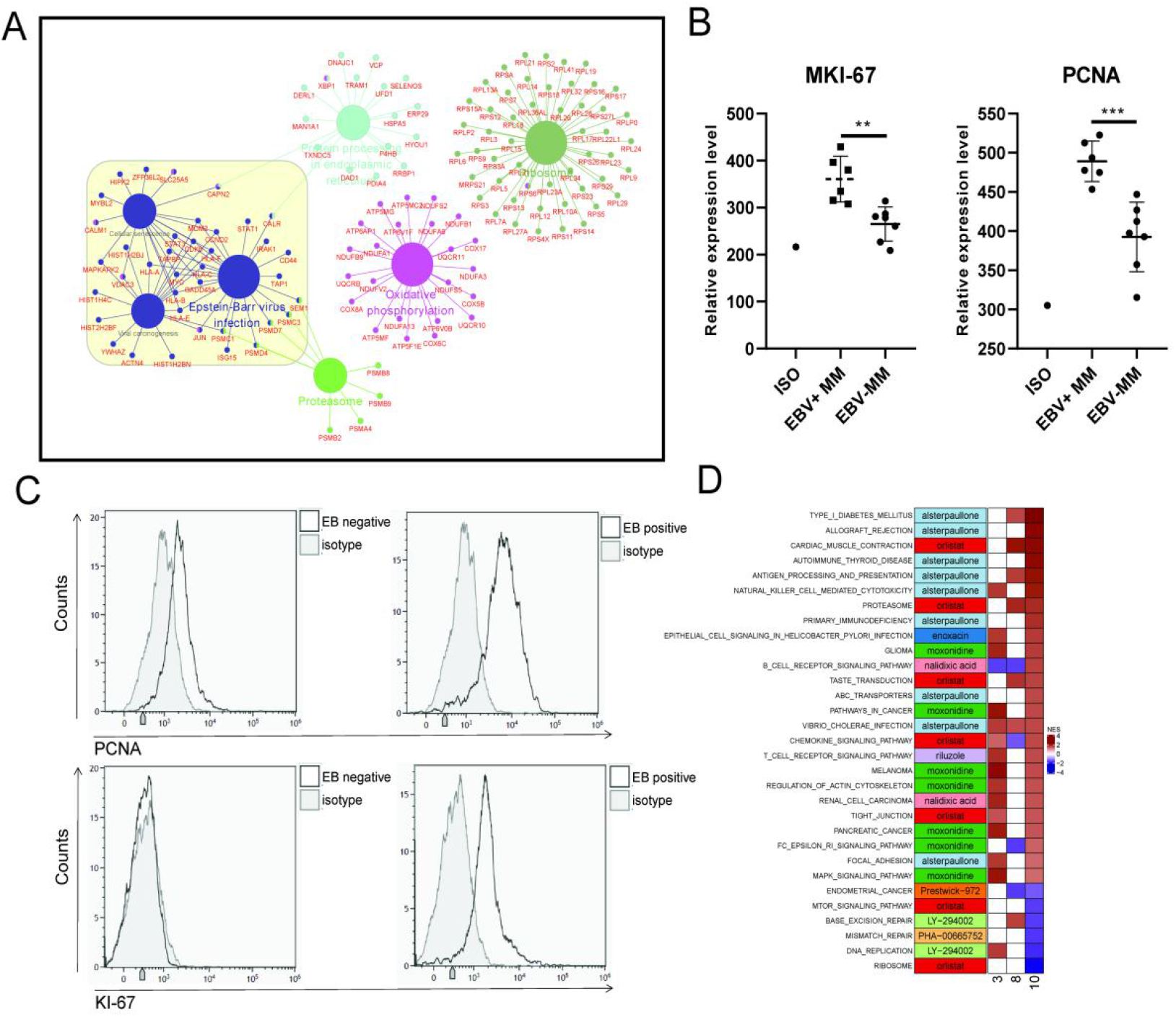
Proliferating plasma cells were increased in EBV positive MM patients. A) DEGs in MM RISS III stage with I stage were obtained and conferred to functional analysis; B) Relative Expression of MKI-67 and PCNA in EBV positive and EBV negative MM patients BM samples; C) Representative FACS peaks of MKI-67 and PCNA in EBV positive and EBV negative MM patients; D) Based on pathways enriched in plasma clusters 3, 8 and 10, potential drug candidates were acquired, and shown with heatmap;

### R-ISS stage-dependent expression analysis highlights functional modules underlying cytotoxic T cell decreases

Accumulating evidence demonstrates that the compromised tumour immune microenvironment (TIME) contributes to MM progression [41], and therapeutic agents targeting the TIME have emerged as promising avenues [13, 42]. As one of the major cytotoxic immune cell types, T cell dysfunction is well acknowledged [43] and immunotherapies such as chimaeric antigen receptor (CAR) T cells [44] and immune checkpoint inhibitors [45] have entered clinical trials. Here, we propose a hypothesis that the proportions of certain cytotoxic T cell populations decrease with MM progression: we stratified them according to the R-ISS system and attempted to identify the functional genes within.

First, re-clustering of T cells generated 21 clusters (**Fig 6A**), T1 to T21, belonging to CD4+ T cells, CD8+ T cells, NK cells and NKT cells (**Fig 6B and 6D**). No biased distribution was observed in 11 samples (**Fig 6C**). The percentages of T1 to T21 in MM versus healthy controls and MM R-ISS I to III are presented in **Fig 6E and Fig 6F**, respectively. It is worth noting that 2 clusters conform to the hypothesis of decreased percentage along with R-ISS stages: T2 and T10. T2 was marked by high expression of CD8A and no expression of NKG7 and was identified as CD8+ T cells. T10 cells express both CD8A and NKG7 and were defined as NKT cells. We concentrate on T2 and T10 in the following work.

**Figure 6.**
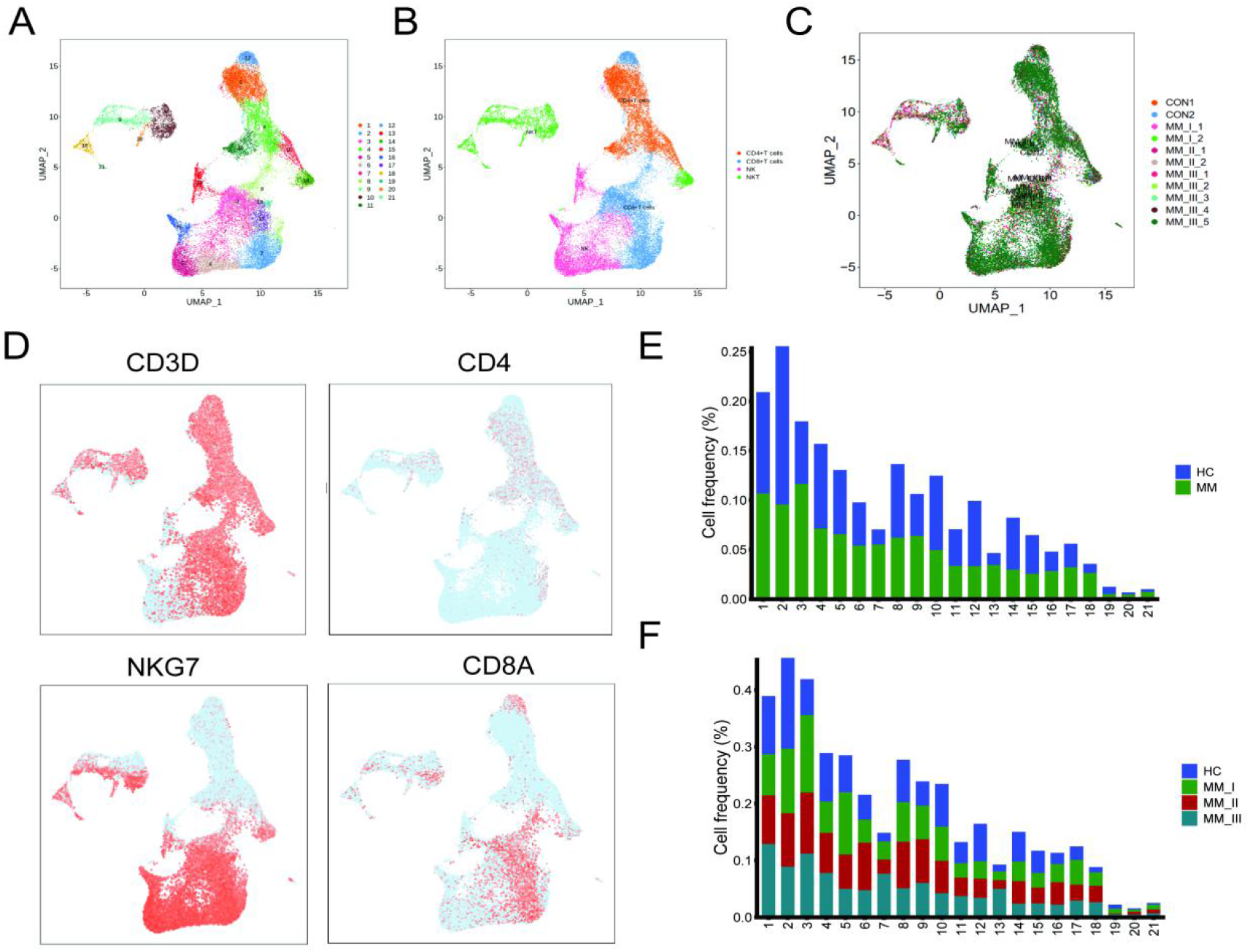
T cell population analysis suggested Stage-dependent CD8+T and NKT cell clusters depletion in MM. A) T cells and NK cells were re-clustered, and 21 clusters were acquired; B) Based on markers expression in D)CD4+T, CD8+T, NK and NKT cells were shown with UMAP; C) UMAP of samples distribution of T cells and NK cells; D) Markers of T cells and NK cells, shown with UMAP; E) Cluster proportions in healthy control and MM groups; F) Cluster proportions in healthy control, RISS-I, II and III MM groups.

To identify R-ISS-dependent gene modules in T2 and T10, the R package MFUZZ [46] was applied. As a result, in T2 CD8+ T cells, 12 gene modules with distinct expression patterns were generated, and module 5 (gradually increased expression with R-ISS stage) and module 3 (gradually decreased expression with R-ISS stage) were chosen for subsequent analysis (**Fig 7A-7B**). As expected, genes in module 5 were functionally related to antigen processing and presentation, T cell activation and haemopoiesis (**Fig 7D**). Surprisingly, genes in module 3 were involved in neutrophil activation (**Fig 7E**), which prompted us to examine neutrophils in MM. Significantly, the proportion of activated neutrophils characterized by CXCR2 expression [47-49] (C5 in Fig 1B) decreased with R-ISS stage (C5 in Fig 1F). Hence, we speculated that the decrease in activated neutrophils among MM R-ISS stages may be attributed to decreased expression of module 3 genes such as CXCR1 [50], ADAM10 [51] and CD47 [52] in T2 cytotoxic cells. For T10, genes in module 1 showed stable expression in healthy controls, R-ISS I and II, while dramatic increases in R-ISS III were observed (**Fig 7C**). Subsequent ClueGo results (**Fig 7F**) revealed genes involved in T cell/lymphocyte differentiation (IL2RA, CD74, CD86, IL7 and RIPK2), adaptive immune response to tumour cells (NECTIN2, IL12A, NRG1 and HSPD1) and nucleotide-binding oligomerization domain (NOD1)-containing signalling pathways (HSPA1A, BIRC3, IRAK1, HSP90AA1 and HSPA1B).

**Figure 7.**
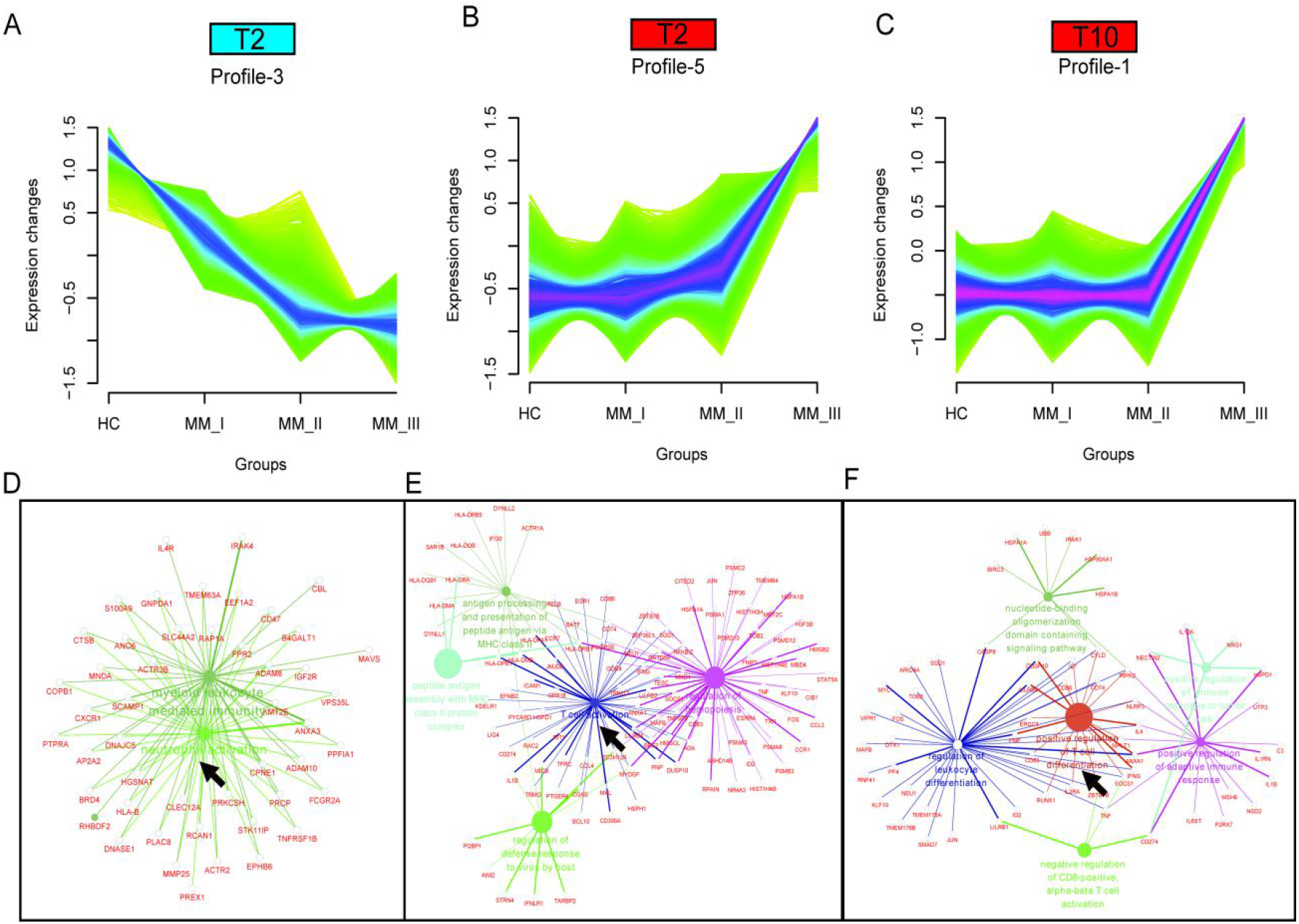
Stage-dependent expression analysis reveals three gene modules in CD8+T and NKT cell clusters. To acquire RISS stage dependent gene expression modules in Cluster 2 and 10 populations, MFUZZ was acquired. Two gene profiles in Cluster2 (T2C3 and T2C5) and Cluster 10 (T10C1) were generated. A) Genes in T2C3 were generally characterized as gradually decreased expression with stage; B) Genes in T2C5 were generally marked with stage dependent increased expression; C) Genes in T10C1 were generally marked stable expression in healthy control, RISS I and II stage, but remarkable elevation in III stage; D) Functional network of genes in T2C3 highlights neutrophil activation of T2 cells; E) Functional network of genes in T2C5 suggested genes in T cell activation; F) Functional network of genes in T10C1 indicates genes in T cell differentiation.

### Ligand-receptor pairs and potential immunotherapeutic targets in CD8+ T-neutrophil and CD8+ T/NKT-plasma cell communication

Finally, we employed CellPhoneDB [53] to interrogate ligand-receptor pairs between T2 CD8+ T cell-C5 myeloid neutrophils (**Fig 8A**) and T2 CD8+ T cell/T10 NKT cell-C10 proliferating plasma cells (**Fig 8C**). As shown in **Fig 8B**, 19 ligand-receptor pairs were proposed as T2 CD8+ T cells to C5 neutrophil modulators, such as chemokine ligand receptors CCL5/CCL3/CCL4L2-CCR1, IFN-IFN receptors (IFNRs), CD99-PILPA and CD52-SIGLEC10. On the other hand, 21 ligand-receptor pairs were proposed as C5 neutrophils to T2 CD8+ T cell modulators, such as chemokine ligand receptors ICAM1/3-SPN/ITGAL, CD55-ADGRE5, SEMA4A-PLXND1, SIRPA-CD44/CD47, IL1B-ADRB2 and CD48-CD244. For T2 CD8+ T cell/T10 NKT cell communication with C10 proliferating plasma cells, common ligand-receptor pairs such as TIGIT-NECTIN3, ADRB2_VEGFB, CD74_COPA, and CD74_MIF were identified. On the other hand, common ligand-receptor pairs such as MDK-SORL1, MIF-TNFRSF14, FAM3C-CLEC2D, and ICAM1-ITGAL were also enriched (**Fig 8D**). Altogether, we found that the ligand-receptor pairs between CD8+ T cells/T10 NKT cells, plasma cells and neutrophils suggested a complex communication network in the MM TIME, which could provide clues for MM progression and therapy.

**Figure 8.**
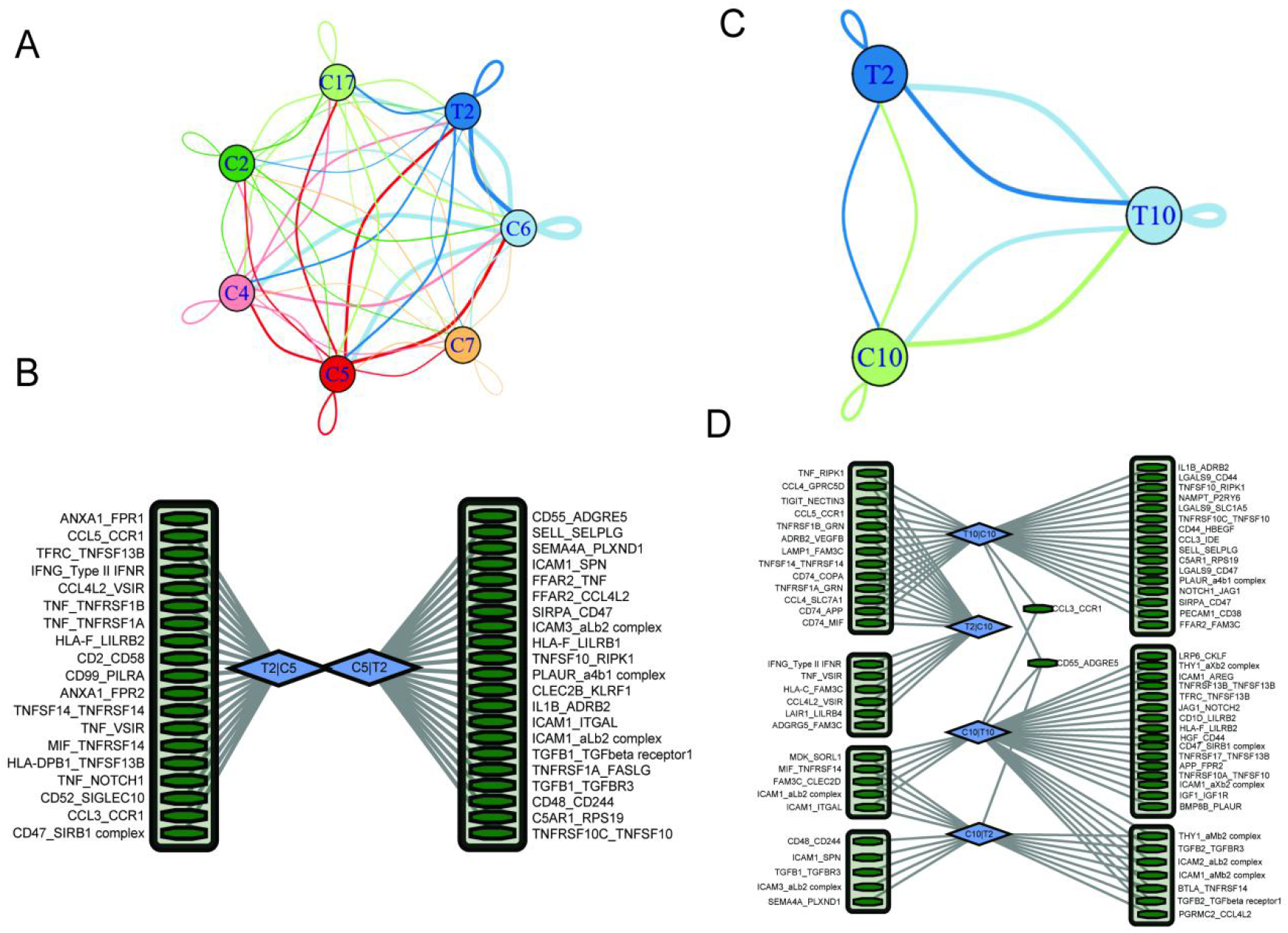
Ligand-receptor pairs and potential immunotherapeutic targets in CD8+T-Neutrophil and CD8+T/NKT-plasma cells communication. A) Cell-cell communications showing the interaction numbers between myeloid cells and T2 CD8+T cells; B) Paracrine ligand-receptor interaction pairs between neutrophils and T2 CD8+T cells; C) Cell-cell communications showing the interaction numbers between C10 plasma cells and T2 CD8+T cells, T10 NKT cells; D) Paracrine ligand-receptor interaction pairs between C10 plasma cells and T2 CD8+T cells, T10 NKT cells;

## Discussion

Multiple myeloma (MM) is characterized by uncontrolled proliferation of monoclonal plasma cells, and the R-ISS System was developed to stratify MM patients into groups I, II and III [2] with distinct outcomes [3] and treatment response [4]. This study identified malignant plasma cells with potent proliferation ability. Moreover, RRM2 and HINT1 with unfavourable prognostic significance in MM were also characterized. RRM2, regulated in a cyclin F-dependent fashion [54], encodes one of two nonidentical subunits for ribonucleotide reductase and is well studied as an oncogene and poor prognostic marker in multiple solid cancer types [55-57]. Meanwhile, RRM2 inhibition induced synergy with gemcitabine in lymphoma [58] and WEE1 inhibitor in H3K36me3-deficient cancers [59]. In MM, RRM2 knockdown alone inhibits MM cell proliferation and induces apoptosis via the Wnt/β-catenin signalling pathway [60]. In contrast, histidine triad nucleotide binding protein 1 (HINT1) encodes a protein that hydrolyses purine nucleotide phosphoramidate substrates and is mostly believed to be a tumour suppressor in multiple cancer types [61-63]. Another finding in malignant plasma cells is the involvement of genes (MDM2, CCND2, CDK6, STAT3, HLA-F, HLA-B, HLA-C, HLA-E, JUN, PSMC1) in Epstein-Barr virus (EBV) infection and viral carcinogenesis [64-67]. Further in-depth studies of their roles in EBV infection and plasma cell proliferation are needed in the future.

Intriguingly, we also found a rare plasma cell proportion with cytotoxic activities (high expression of NKG7 and GZMA) in the BM microenvironment of MM patients for the first time. T cells and NK cells are the two main types of cytotoxic immune cells in previous studies. Coincidentally, plasma/B cells producing granzyme B (GZMB) also possess cytotoxic activities and induce HCT-116 cell death [27]. To be cautious, we first validated the existence of these cytotoxic NKG7/GZMA+ plasma cells in another single-cell dataset. If possible, we collected more MM samples and applied FACS to verify its existence, although the proportion in all plasma cells was small. Furthermore, to explore the function of NKG7/GZMA+ plasma cells, in vitro cytotoxicity assays should be conducted to study the function of NKG7/GZMA+ plasma cells in inducing cell death in MM. Altogether, this discovery offers an alternative option for the cytotherapy of MM.

In addition, we also identified multiple immunotherapeutic targets in MM. CD24 is a highly expressed, anti-phagocytic signal in several cancers and demonstrates therapeutic potential for CD24 blockade in cancer immunotherapy [68]. ICAM-1 antibodies showed potent anti-myeloma activity in multiple studies [69] [70] [71]. CD44 mediates resistance to lenalidomide in multiple myeloma [72], and CD44-targeted T cells mediate potent anti-tumour effects against multiple myeloma [73]. CD47 on macrophages represents a “do-not-eat-me” immune checkpoint [74]. CD48 was expressed on more than 90% of MM plasma cells, and administration of the anti-CD48 mAb significantly inhibited MM growth [75]. CD74 is predominantly expressed in malignant plasma cells, and anti-CD74 mAbs internalize very rapidly and have shown efficacy in B-lymphoma models [76] [77]. MIF is an important player and a novel therapeutic target in MM. Inhibiting MIF activity will sensitize MM cells to chemotherapy [78]. MIF plays a crucial role in MM sensitivity to PIs and suggests that targeting MIF may be a promising strategy to (re)sensitize MM to treatment [79]. T-cell immunoglobulin and ITIM domains (TIGIT) are other immune checkpoint receptors known to negatively regulate T-cell functions. MM progression was associated with high levels of TIGIT expression on CD8 T cells. TIGIT CD8 T cells from MM patients exhibited a dysfunctional phenotype characterized by decreased proliferation and inability to produce cytokines in response to anti-CD3/CD28/CD2 or myeloma antigen stimulation. TIGIT immune checkpoint blockade restores CD8 T-cell immunity against multiple myeloma [80]. Nectin-2 expression on malignant plasma cells is associated with better response to TIGIT blockade in multiple myeloma [81]. Myeloma escape after stem cell transplantation is a consequence of T-cell exhaustion and is prevented by TIGIT blockade [82].

In conclusion, we constructed a single-cell transcriptome atlas of bone marrow in normal and R-ISS-staged MM patients. Focusing on PCs, we identified and validated the existence of GZMA+ cytotoxic PCs. In addition, a malignant PC population with high proliferation capability (proliferating PCs) was clinically associated with EBV infection and unfavourable prognosis.

Ribonucleotide Reductase Regulatory Subunit M2 (RRM2), a specific marker of proliferating PCs, was shown to induce MM cell line proliferation and serve as a detrimental marker in MM. Subsequently, three R-ISS-dependent gene modules in cytotoxic CD8+ T and NKT cells were identified and functionally analysed. Finally, cell-cell communication between neutrophils and proliferating PCs with cytotoxic CD8+ T and NKT cells was investigated, which identified intercellular ligand receptors and potential immunotargets such as SIRPA-CD47 and TIGIT-NECTIN3. Collectively, the results of this study provide an R-ISS-related single-cell MM atlas and reveal the clinical significance of two PC clusters, as well as potential immunotargets in MM progression.

## Acknowledgements

We thank all the patients and their families for participating in this study. The present study was supported by the grant from the National Natural Science Foundation of China (82002212, 81870683, 82070928), the Science & Technology Department of Sichuan Province (19YJ0593, 2020ZYD035, 2020YJ0460, and 2020JDTD0028), Department of Sichuan Provincial Health (19PJ117), the Sichuan Provincial People’s Hospital (2018LY03), the Chengdu Science and Technology Bureau (2019-YF05-00572-SN), the China Postdoctoral Science Foundation Grant (2019M663567), the fundation of Basic scientific research in Central Universities of University of Electronic Science and technology (ZYGX2020J024).

## Authors’ contributions

Bo Gong, Ling Zhong and Yunbin Zhang conceived the idea. Ling Zhong, Xinwei Yuan, JiangTao, Huan Li, and Jialing Xiao collected the specimen and prepared single-cell suspension for sequencing. Ling Zhong, Qian Zhang, Xinwei Yuan, Ping Shuai, Liang Wang, Yuping Liu, Man Yu and Yi Shi finished the bioinformatics analysis. Lan Luo and Chenglong Li accomplished the flow cytometry. Jialing Xiao finished immunofluorescence staining. Ping Shuai, and Yuping Liu finished the qPCR. Ling Zhong, Bo Gong, and Yunbin Zhang wrote the manuscript. All authors reviewed and approved the manuscript.

## Data availability

Sequencing data have been deposited in GEO under accession code GSE176131.

## Material and methods

### Patients and sample collection

This study included 9 MM patients diagnosed as active MM according to International Myeloma Working Group guideline, and two age-matched normal control (transplant donor). For FACS analysis of MKI67 and PCNA in plasma cells, samples from 3 healthy donors, 5 RISS-I, 4 R-ISS-II and 6 R-ISS III were collected. Written informed consents were obtained from all subjects. All experimental procedures were approved by the Institutional Review Board of Sichuan Provincial People’s Hospital and carried out in accordance with the principles of the Declaration of Helsinki.

Bone marrow (BM) aspirates were collected into EDTA-containing tubes, and lysed using Versalyse Lysing Solution (cat. no. A09777; Beckman Coulter, Inc.). Mononuclear cells were isolated using a Ficoll gradient (density 1.077 g/ml,cat. no. 07801; STEMCELL Technologies). Fresh single-cell suspensions were used for ScRNA seq. Aliquots of the same bone biopsy were analyzed by fluorescence in situ hybridization (FISH) and multi-parameter flow cytometry (MFC, Navios Beckman Coulter, Inc.) as parts of the routine clinical diagnosis. In MFC, cell populations were considered abnormal if they have an atypical differentiation pattern, an increased or decreased expression level of normal antigens, an asynchronous maturational pattern or express aberrant antigens [83].

### ScRNA-Seq library construction and sequencing

Single-cell RNA-Seq libraries were prepared with Chromium Single cell 3’ Reagent v3 Kits according to the manufacturer’s protocol. Single-cell suspensions were loaded on the Chromium Single Cell Controller Instrument (10×Genomics) to generate single cell gel beads in emulsions (GEMs). Briefly, about 2×10^5^PBMC single cells were suspended in calcium- and magnesium-free PBS containing 0.04% weight/volume BSA. About 22,000 cells were added to each channel with a targeted cell recovery estimate of 10,000 cells. After generation of GEMs, reverse transcription reactions were engaged barcoded full-length cDNA followed by the disruption of emulsions using the recovery agent and cDNA clean up with DynaBeads® MyOne™ Silane Beads (Thermo Fisher Scientific). cDNA was then amplified by PCR with appropriate cycles which depend on the recovery cells. Subsequently, the amplified cDNA was fragmented, end-repaired, A-tailed, index adaptor ligated and library amplification. Then these libraries were sequenced on the Illumina sequencing platform (NovaSeq6000) and 150 bp paired-end reads were generated. The GEM generation, library construction and sequencing were performed by OE Biotech CO., LTD (Shanghai, China).

### ScRNA-Seq data processing

The Cell Ranger software pipeline (version 3.1.0) provided by 10×Genomics was used to demultiplex cellular barcodes, map reads to the genome and transcriptome using the STAR aligner, and down-sample reads as required to generate normalized aggregate data across samples, producing a matrix of gene counts versus cells. We processed the unique molecular identifier (UMI) count matrix using the R package Seurat[84] (version 3.1.1). To remove low quality cells and likely multiplet captures, which is a major concern in microdroplet-based experiments, we applied criteria to filter out cells with UMI/gene numbers out of the limit of mean value ± 2 fold of standard deviations assuming a Guassian distribution of each cells’ UMI/gene numbers. Following visual inspection of the distribution of cells by the fraction of mitochondrial genes expressed, we further discarded low-quality cells where >20% of the counts belonged to mitochondrial genes. After applying these QC criteria, 103043 single cells were retained for downstream analyses. Library size normalization was performed with Normalize Data function in Seurat[84] to obtain the normalized count. Specifically, the global-scaling normalization method “LogNormalize” normalized the gene expression measurements for each cell by the total expression, multiplied by a scaling factor (10,000 by default), and the results were logtransformed.

Top variable genes across single cells were identified using the method described in Macosko *et al*[85]. The most variable genes were selected using FindVariableGenes function(mean.function=ExpMean, dispersion.function= LogVMR) in Seurat[84]. To remove the batch effects in single-cell RNA-sequencing data, the mutual nearest neighbors (MNN) presented by Haghverdi *et al* was performed with the R package batchelor[84].Graph-based clustering was performed to cluster cells according to their gene expression profile using the FindClusters function in Seurat[84]. Cells were visualized using a 2-dimensional t-distributed stochastic neighbor embedding (t-SNE) algorithm with the RunTSNE function in Seurat[84]. We used the FindAllMarkers function(test.use = bimod) in Seurat[84] to identify marker genes of each cluster. For a given cluster, FindAllMarkers identified positive markers compared with all other cells. Then, we used the R package SingleR[86], a novel computational method for unbiased cell type recognition of scRNA-seq, with the reference transcriptomic datasets ‘Human Primary Cell Atlas’[87] to infer the cell of origin of each of the single cells independently and identify cell types.

Differentially expressed genes (DEGs) were identified using the FindMarkers function (test.use = MAST) in Seurat[84]. P value < 0.05 and |log_2_foldchange| > 0.58 was set as the threshold for significantly differential expression. GO enrichment and KEGG pathway enrichment analysis of DEGs were respectively performed using R based on the hypergeometric distribution.

### Large-scale chromosomal copy number variation analysis

The normalized scRNA-seq gene expression matrices were used to estimate CNV profiles with inferCNV R package[88]. Genes were sorted based on their chromosomal location and a moving average of gene expression was calculated using a window size of 101 genes. The expression was then centered to zero by subtracting the mean. The de-noising was carried out to generate the final CNV profiles.

### Cell cycle analysis

Cell cycle genes were defined as those with a “cell cycle process” Gene Ontology annotation (downloaded from MSigDB version 3.1). We defined four cell cycle signatures (G1/S, S, G2/M, and M) as the average expression [log2(TPM + 1)] of phase-specific subsets of the cell cycle genes. We refined these signatures by averaging only over those genes whose expression pattern in our data correlated highly (r > 0.5) with the average signature of the respective cell cycle phase (before excluding any gene) in order to remove the influence of genes.

### Cell culture

Human normal plasma cells were separated from the peripheral blood of healthy donors using flow cytometry. All donors were healthy volunteers who had not previously received any drugs associated with immunological diseases. Briefly, peripheral blood mononuclear cells were separated by Ficoll®-Hypaque centrifugation from peripheral blood. Plasma cells were isolated from mononuclear blood cells using CD138 microbeads (Miltenyi Biotec, Inc.) according to the manufacturer’s instructions [89]. The isolated plasma cells were then cultured. The U266 and MM1.S cell lines were purchased from the American Type Culture Collection. RPMI-1640 medium (Gibco; Thermo Fisher Scientific, Inc.) containing 10% fetal bovine serum (Gibco; Thermo Fisher Scientific, Inc.) and 1% dual antibiotics (penicillin 100 U/ml; streptomycin 0.1 mg/m1; Sigma-Aldrich; Merck KGaA) was used to culture all cells at 37°C with 5% CO2.

### RRM2 and HINT1 silencing with shRNAs

Short hairpin RNAs specifically targeting RRM2 and HINT1 were designed and synthesized by. RRM2 shRNAs were shRNA1 (5’-3’): GGAGCGATTTAGCCAAGAA, shRNA2 (5’-3’): GCCTCACATTTTCTAATGA and shRNA3 (5’-3’): GAAAGACTAACTTCTTTGA. HINT1 shRNAs were: shRNA1 (5’-3’): GGTGGTGAATGAAGGTTCA, shRNA2 (5’-3’): GTGATACCCAAGAAACATA and shRNA3 (5’-3’): GTCTGTCTATCACGTTCAT.

The shRNAs were cloned into pcDNA3.1, and transfected into the MM cells. Transfection was performed according to the manufacturer’s protocol using Lipofectamine® 3000 reagent. Following co-culture for 12 h at 37°C, the medium was replaced with culture media and the transfected cells were used for subsequent experiments. Forty-eight hours post transfection, RRM2 and HINT1 expression in the cells was determined by reverse transcription quantitative polymerase chain reaction (RT-qPCR), and the transfection efficiency was verified.

#### RNA extraction and reverse transcription-quantitative RT-qPCR

Total RNA from cells was extracted using TRIzol (Invitrogen; Thermo Fisher Scientific, Inc.). cDNA was synthesized using the PrimeScript RT reagent kit (Takara Biotechnology Co., Ltd.) at 25°C for 5 min, 37°C for 30 min and 85°C for 5 sec. qPCR analysis was performed using the StepOnePlus Real-Time PCR system (Applied Biosystems; Thermo Fisher Scientific, Inc.) with SYBR Premix EX Taq kit (Takara Biotechnology Co., Ltd.). Primer sequences for RRM2, HINT1were as following: RRM2 Forward 5’-3’: CCAATGAGCTTCACAGGCAA, Reverse 5’-3’: TGGCTCAAGAAACGAGGACT. HINT1Forward5’-3’:TTGCCGACCTCCAAGAACAT, Reverse 5’-3’: CCCTCAAGCACCAACACATT. Relative expression was calculated using the 2−ΔΔCq method [90].

#### CCK8 assays for cell proliferation

Cell proliferation assay was performed with Cell Counting Kit-8 (Dojindo, Kumamoto, Japan) according to the manufacturer’s instruction. Twenty-four hour after transfection, cells were seed in 96-well plates at 1 × 104 U266 cells per well. The proliferative ability of U266 cells was determined at 0, 24 and 48 h. The absorbance was measured at 450 nm using a microplate spectrophotometer (Molecular Devices, Sunnyvale, USA).

### Statistical analysis

Statistical analysis and graph representations were performed using SPSS v.13.0 software (SPSS Inc., Chicago, IL) and GraphPad Prism 8 Software (GraphPad, San Diego, CA), respectively. One-way ANOVA with post hoc Tukey’s test was used to compare differences between multiple groups. For cell assays, data are presented as the mean ± standard deviation (SD) and were compared using either Student’s *t* test or the Mann-Whitney *U* test. The Kaplan-Meier method was used for survival analyses. P<0.05 was considered statistically significant.

